# A human-specific genetic modifier reconfigures large-scale cortical network dynamics underlying behavioral performance

**DOI:** 10.64898/2026.06.24.734394

**Authors:** Hanzhi T. Zhao, Taylor R. Anderson, Ewoud R. E. Schmidt

## Abstract

Cortical expansion in the human lineage was accompanied by alterations in cortical circuit architecture, including an elaboration of cortico-cortical connectivity. Yet how these changes reshape network computation to support behavior remains poorly understood. Here we leverage a mouse model expressing *SRGAP2C*, a human-specific gene duplication that modifies cortical circuit development and increases cortico-cortical connectivity, to ask how this remodeled architecture shapes network dynamics during learning. As mice acquire expertise on a sensory discrimination task, *SRGAP2C* drives broader interhemispheric correlation, distributed task-relevant encoding, and enhanced directed influence from frontal and associative regions. Texture representations in primary sensory cortex become more separable during motor preparation, supporting improved discrimination under demanding conditions. These findings demonstrate that human-specific changes in cortical circuit architecture do not simply scale up but rather reconfigure the cortical dynamics that organize sensory-to-motor transformation in a manner that tracks behavioral performance under demanding conditions, linking genomic innovations in the *Homo* lineage to large-scale cortical network function.

## INTRODUCTION

The neocortex is organized as a densely interconnected network of functionally specialized areas, and the large-scale pattern of cortico-cortical connectivity is a fundamental determinant of how sensory information is processed, transformed, and ultimately used to guide behavior^1–5^. Across mammalian evolution, expansion of the neocortex has been accompanied by a parallel elaboration of cortico-cortical projections, with a selective increase in cortico-cortical projection neurons in the human lineage^6–10^, suggesting that increased inter-areal connectivity may be a key substrate of enhanced cognitive capacity. Yet the relationship between cortical circuit architecture and information processing remains poorly understood, leaving unknown both how the elaboration of cortico-cortical connectivity that accompanied cortical expansion shaped network-level computation, and whether the connectivity innovations that distinguish the human cortex impact how cortical networks process information and support behavior. Because the connectivity changes that accompanied cortical evolution were ultimately driven by changes in the genome, bridging this gap requires approaches that can link a genetically defined change in cortico-cortical connectivity to the dynamics of large-scale cortical networks during learning and task performance.

*SRGAP2C* arose approximately 2-3 million years ago in the ancestral genome of the *Homo* lineage^11,12^. *SRGAP2C* encodes a truncated protein that binds to and inhibits the function of its ancestral paralog SRGAP2A, a synaptic regulator conserved across mammals, including mice and humans. SRGAP2A functions to limit synaptic density and promote spine maturation in cortical pyramidal neurons^12–15^. This inhibition leads to an increase in the density of both excitatory and inhibitory synapses received by cortical pyramidal neurons and protracted spine maturation, features that recapitulate traits characterizing human cortical neurons^12,13,16,17^. Critically, the increased cortical connectivity originates selectively from local and long-range cortico-cortical projections, with no change in subcortical inputs^18^. Our previous work using a mouse model humanized for *SRGAP2C* expression demonstrated modified sensory-evoked responses (in a passive, non-learned context) and improved learning on a cortex-dependent texture discrimination task^18^, establishing that *SRGAP2C*-induced changes in cortical circuit architecture has measurable functional consequences. Yet, how this remodeled architecture reshapes the large-scale network dynamics that are engaged as animals learn and perform the task, i.e., whether cortical areas communicate differently, whether sensory representations are reorganized at the population level, and whether the architecture of task-related activity is fundamentally altered or merely scaled, has remained unaddressed. Answering these questions requires moving beyond single-area measurements to characterize the distributed, time-resolved network dynamics that unfold as animals become expert performers.

Here we use widefield and two-photon calcium imaging, generalized linear modeling, Granger causality analysis, and psychometric behavioral testing to ask how *SRGAP2C*-mediated connectivity shapes cortical network dynamics during performance of a sensory discrimination task. We find that in *SRGAP2C* mice cortical networks adopt distinct organizational strategies as they become expert learners. *SRGAP2C* mice develop broader spatial correlation, a distributed encoding architecture during motor preparation, and enhanced directed influence from top-down cortical regions across a bilateral cortical network. At single-cell resolution, stimulus representations become more separable during the preparatory epoch in *SRGAP2C* mice. Importantly, these changes in directed influence and population geometry are correlated with improved discrimination performance when stimuli become more similar. These findings demonstrate that the shaping of cortical connectivity by the human-specific gene *SRGAP2C*, including an increase in long-range cortico-cortical connectivity, does not simply scale existing network dynamics. Instead, it reconfigures how the cortex organizes the transformation of sensory evidence into motor preparation in a manner that tracks behavioral performance under demanding conditions, linking genomic innovations in the *Homo* lineage to large-scale cortical function.

## RESULTS

### Equivalent task acquisition in WT and *SRGAP2C* mice provides a uniform baseline for interrogating cortical dynamics

To investigate how *SRGAP2C*-mediated reconfiguration of cortical architecture shapes network dynamics that underlie the learning advantage of *SRGAP2C* mice, we used the mouse whisker system as a model. This system is well-suited for this purpose: it supports cortically dependent learned discriminations that can be parametrically varied in difficulty in order to probe the limits of sensory processing^19–21^, and relies on a well-characterized pathway from sensory periphery through barrel cortex with defined cortico-cortical projections to motor and association areas^22,23^. Beyond its utility as a tractable model system, the barrel cortex circuit shares the canonical cortico-cortical architecture, including layered pyramidal neurons, defined long-range projections to motor and association areas, and feedforward and feedback pathways, that organizes sensory processing across mammalian neocortex^24,25^. The principles governing how cortical connectivity shapes sensory processing and learning in this circuit may therefore generalize beyond the somatosensory system, providing insights relevant to understanding how connectivity innovations in the human cortex may shape cognition more broadly.

We trained WT and conditional *SRGAP2C*-expressing mice, in which *SRGAP2C* is selectively expressed in cortical excitatory neurons via a Cre-dependent knock-in allele (NexCre^26^-*Rosa26^LSL-SRGAP2C-HA^),* referred to hereafter as *SRGAP2C* mice^18^, on a two-alternative forced choice (2AFC) whisker-based texture discrimination task (**Fig. 1a**). Head-fixed mice discriminated between rough and smooth textures and reported their choice with a directed lick to the corresponding port for a water reward. A brief delay was enforced between texture presentation and opening of the response window, as indicated by a visual light cue, to temporally dissociate sensory processing from motor execution for our imaging strategies. Premature licks during the sensing or delay period resulted in immediate trial termination and a timeout period, discouraging impulsive responding (**Fig. 1a**).

**Fig. 1.**
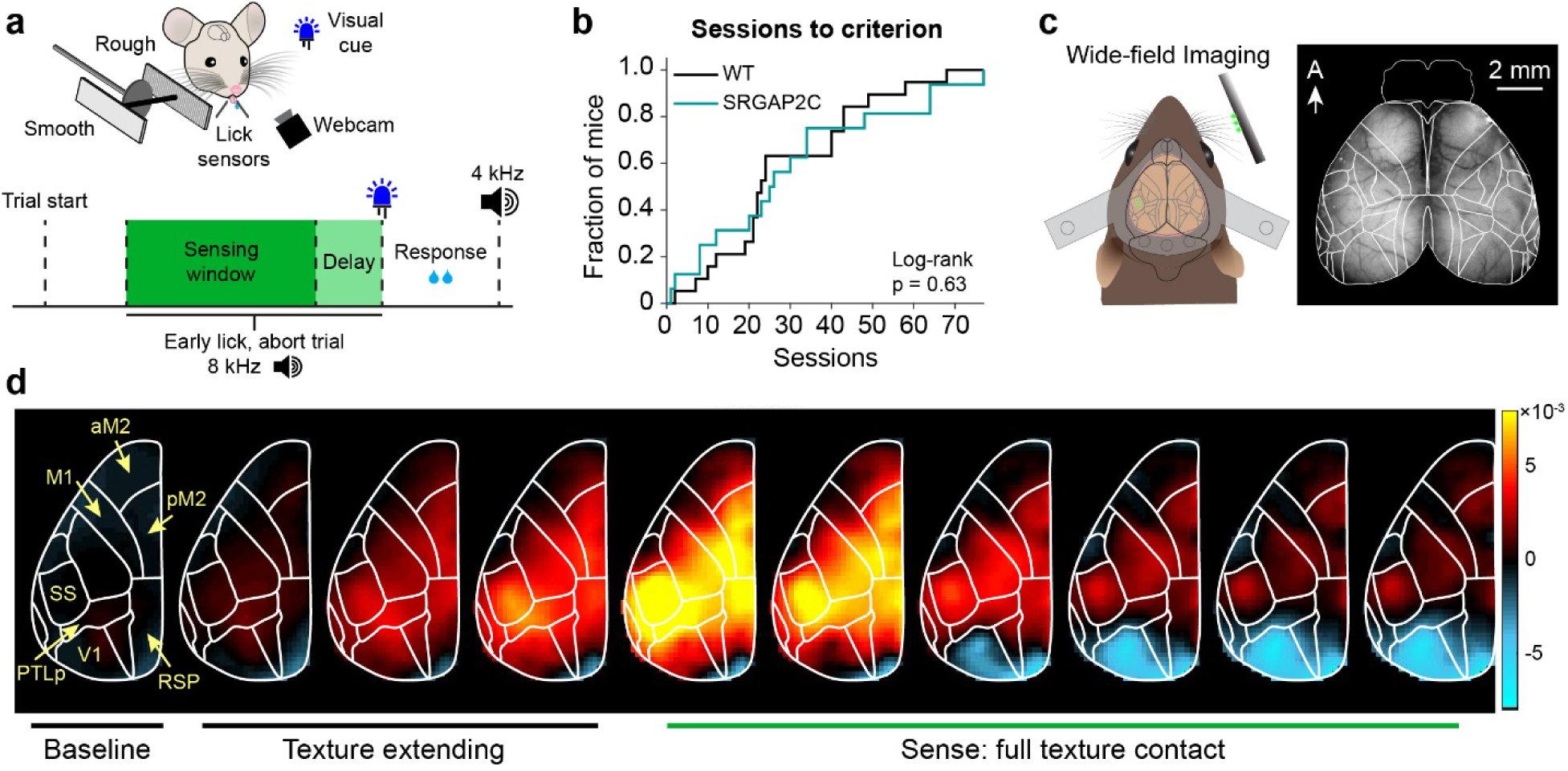
Widefield imaging during a two-alternative forced choice texture discrimination task. **a**, Top, schematic of the behavior task setup. Bottom, schematic of trial structure. **b**, Cumulative histogram showing the number of sessions needed to reach expert performance. P = 0.63 log-rank (*n* = 19 WT mice and *n* = 16 *SRGAP2C* mice). **c**, Left, schematic of the widefield imaging field of view. Right, representative fluorescence image. **d**, Sequential cortical fluorescence maps showing mean ΔF/F activity across the dorsal cortex for a representative animal, registered to the Allen Common Coordinate Framework (CCF) with region outlines overlaid. Warmer colors indicate increases and cool colors indicate decreases in ΔF/F relative to baseline. Regions of interest used in subsequent analyses are labeled.

A key challenge in interpreting genotype differences in cortical dynamics is disambiguating effects on neural processing from effects that are secondary to differences in behavioral performance. Our previous work demonstrated a performance advantage in *SRGAP2C* mice on demanding texture discriminations^18^, raising the possibility that any observed differences in cortical activity could reflect differences in behavioral performance level rather than circuit-level processing per se. To isolate genotype differences in cortical circuit dynamics from differences in behavioral performance, we trained mice on a smooth versus rough texture pair, a version of the task that is sufficiently easy to support equivalent performance across genotypes, in contrast to the more demanding rough-rough discrimination on which *SRGAP2C* mice show a performance advantage^18^ (also see **Fig. 5** for improved performance of *SRGAP2C* mice on more demanding discriminations). Both genotypes successfully acquired the task, and learning curves revealed no significant difference in acquisition rate (**Fig. 1b** and **Supplementary Fig. 1a, b**). Throughout training, response bias and early lick rates were comparable across genotypes, indicating similar levels of task engagement (**Supplementary Fig. 1b**). Response latency was measured separately for correct and incorrect trials. While incorrect trials showed modestly longer latencies as expected, no genotype differences were observed for either outcome (**Supplementary Fig. 1c**). Whisker and body movement during trials did not differ between genotypes (**Supplementary Fig. 1d**).

These results confirm that on the smooth versus rough discrimination paradigm, *SRGAP2C* and WT mice learn and perform at equivalent levels across all measured behavioral dimensions, establishing a uniform performance baseline necessary for interpreting subsequent differences in cortical dynamics.

### Learning-dependent expansion of S1 functional connectivity in *SRGAP2C* mice

Because *SRGAP2C* alters cortical circuit architecture, including enhanced local and long-range cortico-cortical inputs to cortical pyramidal neurons^18^, its effects on cortical activity during task performance are unlikely to be confined to a single area. To characterize how this altered architecture shapes cortical dynamics at the level of distributed inter-areal networks, we performed longitudinal widefield calcium imaging^27,28^ of the dorsal cortex in a subset of WT and *SRGAP2C* mice expressing GCaMP6f (Thy1-GCaMP6f^29^) while they performed the texture discrimination task (**Fig. 1c, d**). We examined the peri-stimulus period, a 2-sec time window around the moment of full texture contact, to evaluate how sensory input arriving in the barrel cortex of primary somatosensory cortex (S1) impacts S1 activity and propagates across the cortex to higher order cortical regions (**Fig. 1d** and **2a**). No genotype differences were observed in this gross measure of cortical dynamics, suggesting that the overall trajectory of task-related cortical reorganization is preserved in *SRGAP2C* mice.

**Fig. 2.**
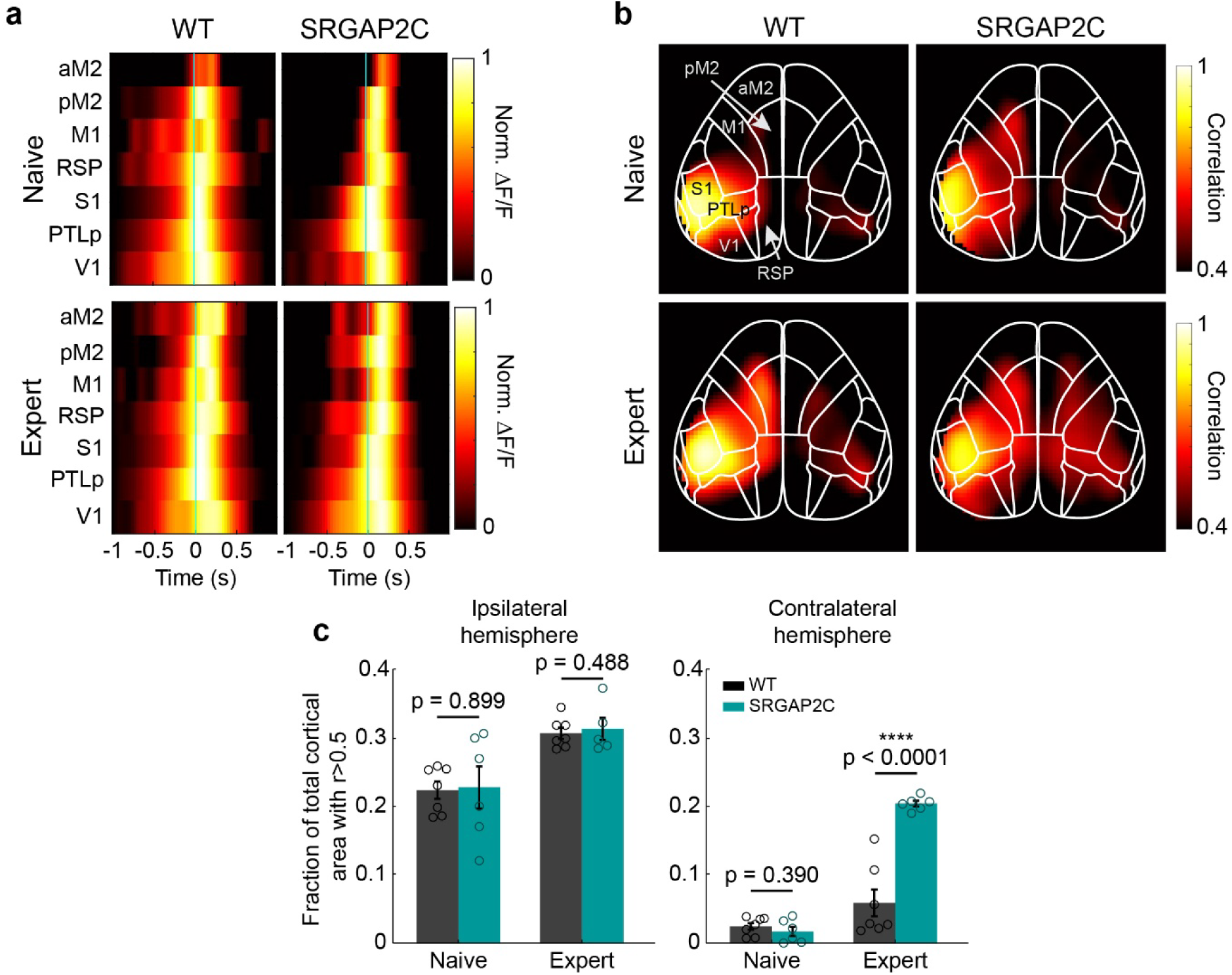
Cortical response dynamics and learning-dependent expansion of S1 correlated activity. **a,** Kymographs of mean ΔF/F for each cortical region of interest across the peri-stimulus window (±1 s around full texture contact) in naïve and expert WT and *SRGAP2C* mice. Green lines denote moment of full texture contact. **b**, Pixelwise Pearson correlation maps seeded to S1 for naïve and expert WT and *SRGAP2C* mice. **c**, Fraction of cortical area with r > 0.5 for the ipsilateral (left) and contralateral (right) hemispheres. ****p < 0.0001, two-sided Mann-Whitney test. Bar graphs plotted as the mean ± s.e.m. Open circle indicate individual animals (*n* = 7 WT, *n* = 6 *SRGAP2C* mice).

To assess whether *SRGAP2C* expression alters the spatial structure of cortical activity during sensory processing, we computed pixel-wise correlation between S1 activity and activity across the dorsal cortex (**Fig. 2b**). In WT mice, we observed strong correlation between S1 and multiple other regions primarily on the ipsilateral side (ipsilateral refers to the hemisphere receiving direct thalamic sensory input, contralateral to the stimulated whiskers), including PTLp, M1, anterior M2, and RSP, with minimal contralateral correlation. Ipsilateral correlation structure was largely similar between groups, although ipsilateral coupling in WT mice is already strong, and a ceiling effect may limit the detection of any further increase in *SRGAP2C* mice. By contrast, *SRGAP2C* mice showed markedly expanded correlation in the contralateral hemisphere, where WT baseline correlations are weaker and the dynamic range for detecting genotype differences is greater (**Fig. 2b, c**). This expanded correlation pattern suggests that sensory-evoked activity propagates more broadly across the dorsal cortex in *SRGAP2C* mice during texture contact. Notably, this difference was absent at the naive stage, emerging only as animals acquired task expertise (**Fig. 2c**). This experience dependence suggests that the expanded cortico-cortical substrate afforded by *SRGAP2C* requires task-relevant learning to be functionally engaged. Increased structural connectivity may lower the threshold for recruiting broader cortical networks, but the task-relevant activity patterns necessary to drive that recruitment emerge only as animals acquire expertise. Taken together, these results suggest that while both genotypes undergo qualitatively similar cortical learning, *SRGAP2C* mice develop a distinctly broader spatial footprint of somatosensory network engagement with task expertise.

### Task-relevant encoding is more spatially distributed in *SRGAP2C* mice

We next asked whether the broader S1-seeded connectivity observed in expert *SRGAP2C* mice reflected functionally meaningful recruitment of cortical territory, or simply passive co-activation that carries little task-associated information. To address this, we used a generalized linear model (GLM) framework^30,31^, which allowed us to test directly whether activity in correlated regions encodes task-relevant variables (**Fig. 3a-b**). Unlike the correlation analysis, which was restricted to the texture-aligned peri-stimulus epoch due to movement-related signal inflation during licking (see Methods), the GLM framework allowed us to separately interrogate both the sensory and motor preparatory epochs by aligning trials to texture presentation and lick onset respectively, providing a more complete picture of how task-relevant information is spatially and temporally organized across the cortex.

**Fig. 3.**
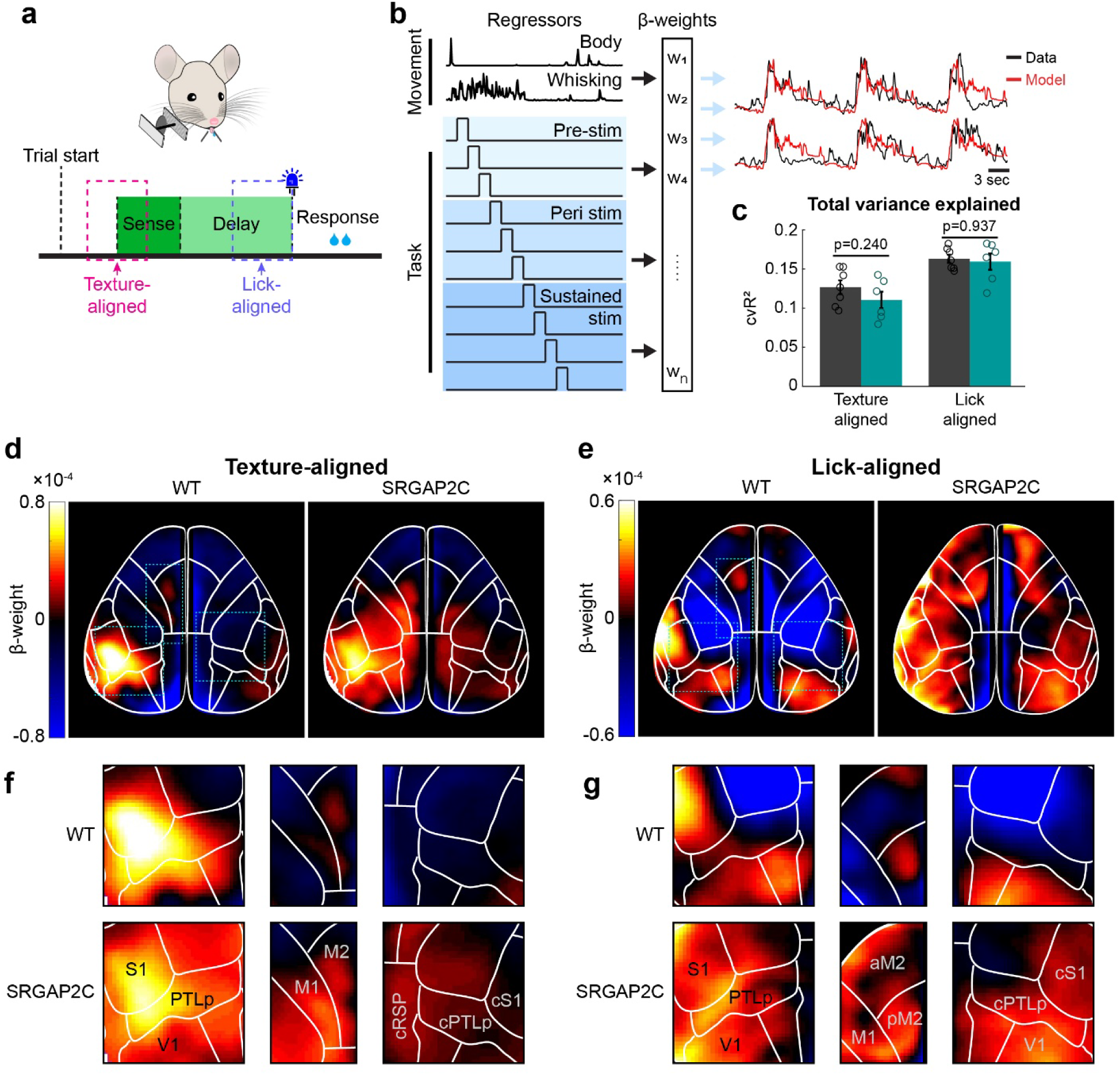
A generalized linear model reveals redistribution of task-related encoding across cortical areas in *SRGAP2C* mice. **a**, Schematic of behavioral task and trial structure indicating the texture-aligned and lick-aligned analysis windows (dashed outlines). **b**, Schematic of the GLM. Movement (top) and task variables (bottom) serve as regressors, with fitted β-weights used to generate predicted fluorescence traces (red) compared against observed data (black). **c**, Total variance explained by the full model for texture-aligned and lick-aligned windows, pairwise two-sample t-test. **d,e**, Spatial maps of mean GLM β-weights during the peri-stimulus epoch for WT and *SRGAP2C* mice, shown for texture-aligned (**d**) and lick-aligned (**e**) windows. **f,g**, Magnified views of selected cortical regions from d and e, respectively. Bar graphs plotted as the mean ± s.e.m. Open circles indicate individual animals (*n* = 7 WT and *n* = 6 *SRGAP2C* mice).

Across both epochs, total variance explained by the full model did not differ between genotypes (**Fig. 3c**), indicating that WT and *SRGAP2C* cortical networks capture equivalent amounts of task-relevant information. However, the spatial distribution of encoding differed markedly. During texture sensing, WT mice showed a focal hotspot of high beta weights centered on S1 with a small region of positive encoding in posterior M2, while *SRGAP2C* mice exhibited more widespread positive beta weights distributed across both hemispheres (**Fig. 3d, f**). A similar pattern emerged during the pre-lick (preparatory) epoch, where WT mice showed localized high beta weights in regions consistent with sensorimotor transformation and motor planning, including primary sensory and frontal cortical regions. Both V1 and contralateral V1 also encoded the task well, likely reflecting the visual cue signaling the opening of the response window. In contrast, *SRGAP2C* mice showed broadly distributed encoding with significant involvement of additional regions including PTLp, M2, cPTLp, cM2, and cS1 (**Fig. 3e, g**). Importantly, the spatial overlap between the core encoding regions of WT mice and the broader pattern in *SRGAP2C* mice suggests that *SRGAP2C* recruits additional cortical territory on top of a preserved sensorimotor encoding framework. Together, these results indicate that the *SRGAP2C*-mediated change in cortical circuit architecture redistributes task-related encoding across a larger cortical territory without altering the total task-related variance explained by the network.

### *SRGAP2C* mice exhibit enhanced separability of texture representations during motor preparation

Widefield imaging revealed that *SRGAP2C* mice develop a broader spatial footprint of cortical network engagement with task expertise, and GLM analysis confirmed that this expanded recruitment reflects encoding of task-related variables rather than passive co-activation. Together these results raise the question of whether this reorganization of large-scale network engagement also reshapes how sensory information is represented at the level of population activity within S1 itself. This question is particularly compelling given that regions showing disproportionate encoding increases in *SRGAP2C* mice, including PTLp and M2, send long-range cortico-cortical projections to S1 that have been shown to play a role in texture discrimination^2,3,22,25,32–41^, and are among the inputs specifically enhanced by *SRGAP2C*^18^. Such inputs are well positioned to shape how sensory information is represented across the population^42,43^. Resolving whether this population-level representation is altered in *SRGAP2C* mice requires single-cell resolution.

To address this, we performed two-photon calcium imaging of layer 2/3 neurons in S1 barrel cortex (**Fig. 4a**) and examined how texture representations are organized in the population activity space of simultaneously recorded neurons, i.e., whether the patterns of activity evoked by rough and smooth textures occupy distinct or overlapping regions of this space. We constructed neural population trajectories for each texture condition in a reduced-dimensional state space and quantified the Euclidean distance between rough and smooth trajectories over time as a measure of this representational separability (**Fig. 4b, c** and **Supplementary Fig. 2**). In the preparatory period preceding the lick response, using population trajectories aligned to lick initiation, *SRGAP2C* mice exhibited significantly greater divergence between rough and smooth trajectories emerging just before the moment of the lick (**Fig. 4c**). This enhanced divergence indicates that by the time of choice and motor execution, population-level representations of the two textures have been pushed farther apart in state space in *SRGAP2C* mice. Examining the distribution of pairwise distances across trials further revealed that while WT mice showed a sharp, narrow peak at short distances, *SRGAP2C* mice exhibited a broader, right-shifted distribution spanning larger pairwise distances (**Fig. 4d**). This shift indicates not only that mean trajectory separation is greater in *SRGAP2C* mice, but that a larger proportion of trial pairs are represented in distinct regions of state space. This result suggests that the broader recruitment of cortical networks in *SRGAP2C* mice coincides with a reorganization of population geometry in S1 during the preparatory epoch.

**Fig. 4.**
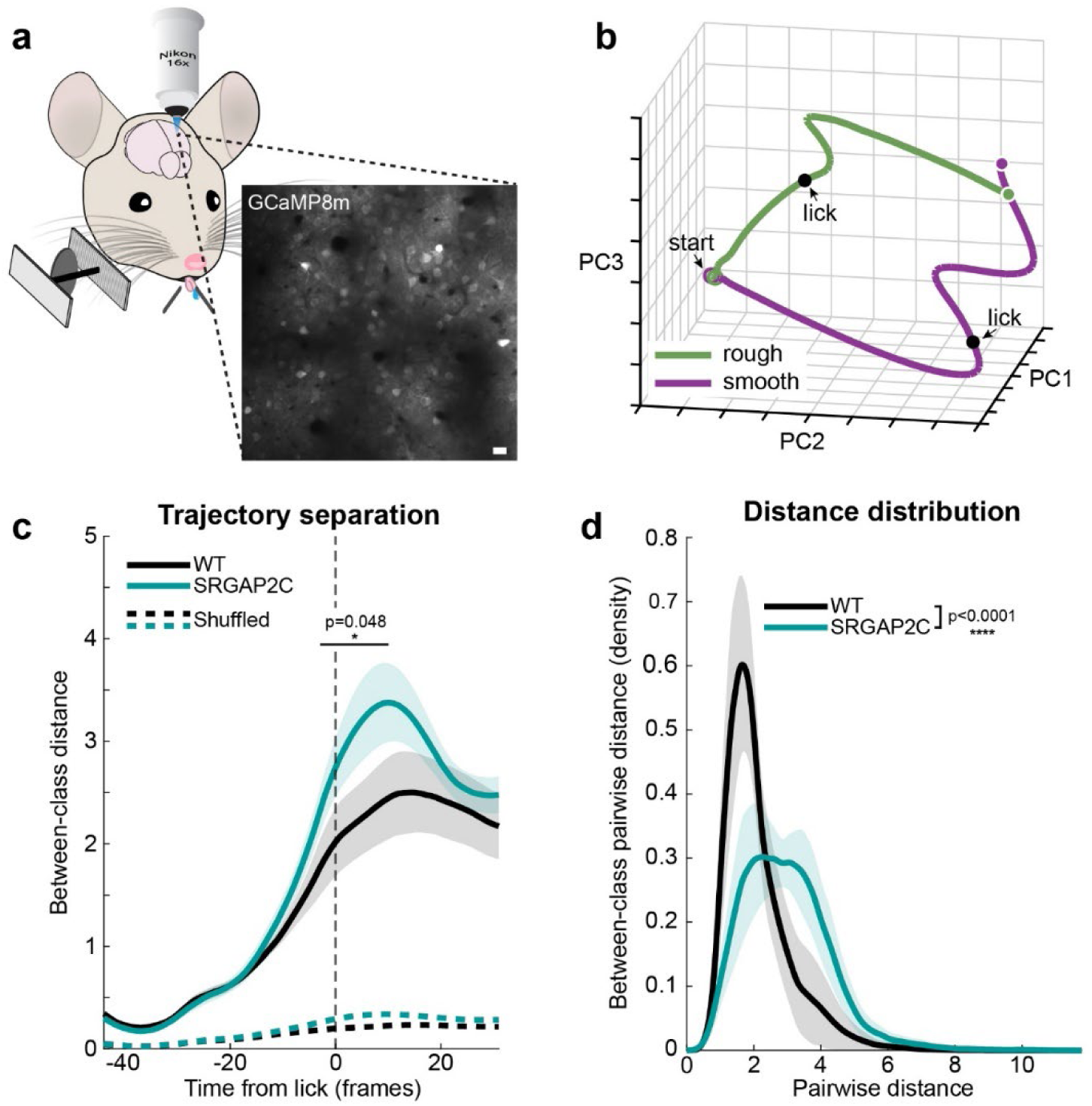
Somatosensory population dynamics reveal enhanced neural trajectory separation in *SRGAP2C* mice. **a**, Representative two-photon field of view. Scale bar, 20 µm **b**, Lick-aligned neural trajectories projected onto the top three principal components (PC1-PC3) for smooth and rough trials in a representative animal. Black dots indicate the moment of lick response. **c**, Euclidean distance between rough and smooth neural trajectories as a function of time relative to lick response for WT and *SRGAP2C* mice shown as mean with shaded areas indicating ± s.e.m., compared to shuffled null distributions (dashed lines). *p<0.05, two-sided Mann-Whitney test. Vertical dashed line indicates lick response. **d**, Distribution of pairwise distances shown as mean ± s.e.m. ****p < 0.0001, Kolmogorov-Smirnov test, *n* = 5 WT and *n* = 7 *SRGAP2C* mice.

### *SRGAP2C* mice maintain discrimination performance under increased task difficulty

Increased distance between population trajectories should have direct implications for the robustness of perceptual decisions. When stimuli representations are farther apart in state space, the boundary between them is more tolerant of noise and perturbations. This property should be especially advantageous when discriminating between stimuli that are more similar, i.e., where population trajectories converge and decision boundaries narrow, predicting that *SRGAP2C* mice should maintain performance better as texture similarity increases.

To directly test whether *SRGAP2C* mice exhibit enhanced performance in distinguishing more similar stimuli, we used a psychometric approach in which we increased texture similarity. We challenged a subset of expert mice from our behavior cohort with a task in which the rough texture was made progressively smoother across three increments, parametrically reducing the contrast between the two stimuli (**Fig. 5a**). Textures are labeled by groove spacing in microns: R200 denotes the standard rough texture used during expert training, while R800, R1400, and R2000 have progressively wider groove spacing and therefore increasingly smooth surface, such that R2000 represents the most similar to the smooth reference and the most demanding discrimination. Performance at expert levels on the standard smooth vs R200 paradigm was comparable between genotypes (**Supplementary Fig. 3a**). To account for inherent variability between mice, we measured the change in performance relative to the expert baseline of each animal on the standard smooth vs. R200 task. WT mice showed an immediate decline in performance (**Fig. 5b**). *SRGAP2C* mice, by contrast, maintained performance near expert baseline levels through the first two texture increments (R800 and R1400), showing minimal impact from the increased difficulty. Only at the most difficult texture combination (smooth vs. R2000) did *SRGAP2C* mice begin to show a measurable decline in performance. To confirm that mice were performing the task based on tactile discrimination rather than alternative sensory cues, we tested all animals on a smooth versus smooth control condition, in which both textures were identical, and no tactile discrimination was possible. Performance dropped to chance levels in both genotypes (**Supplementary Fig. 3b**), confirming that behavior was driven by whisker-based texture discrimination rather than inadvertent use of auditory, visual, or olfactory cues.

**Fig. 5.**
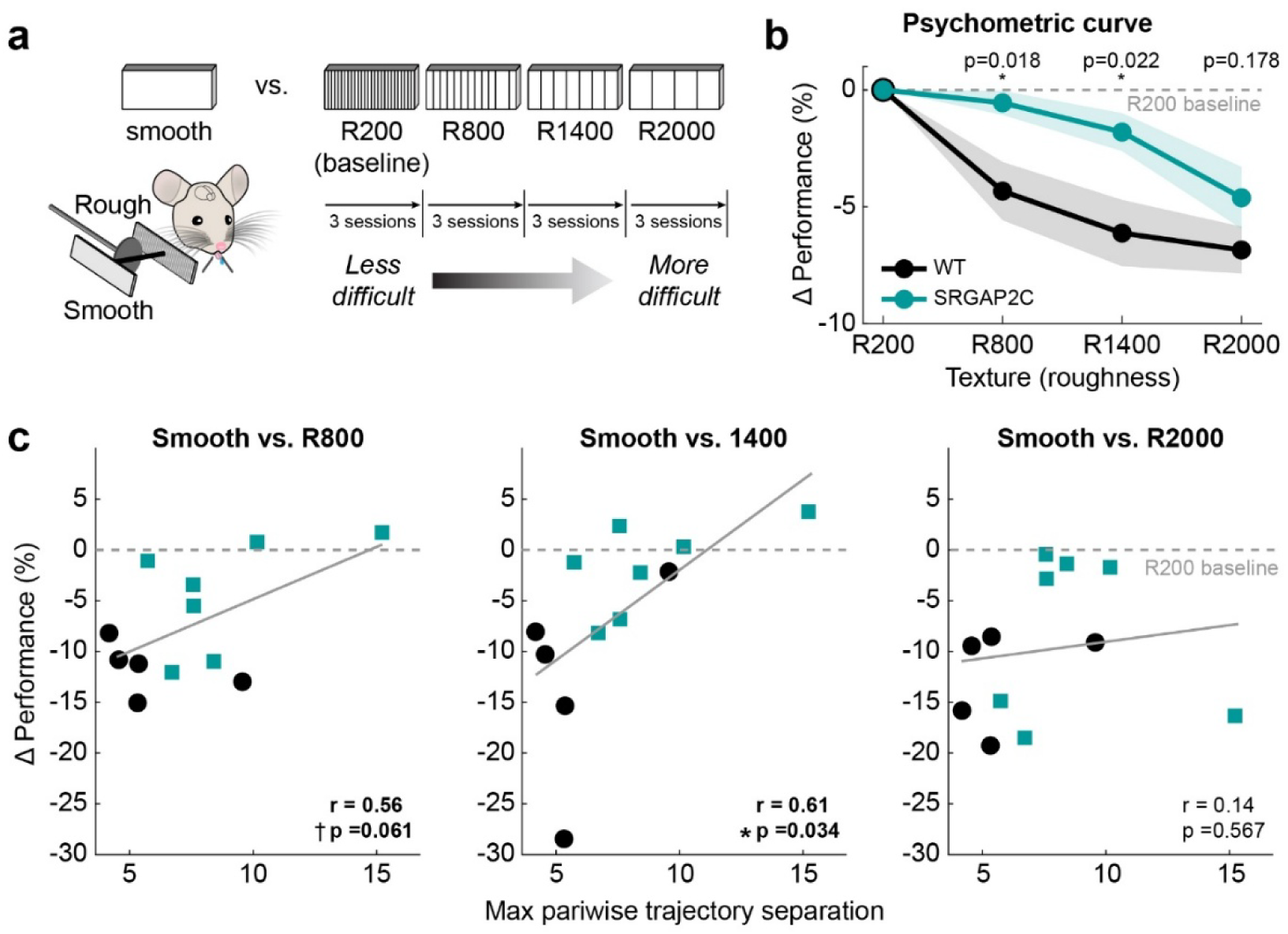
*SRGAP2C* mice maintain superior discrimination performance as texture difficulty increases. **a**, Schematic of the psychometric stimulus series. Textures range from rough (R200) to fine (R2000) with smooth as the reference, presented across three sessions each. **b**, Psychometric curves showing mean performance at each texture relative to the smooth vs. R200 baseline. The shaded area indicates ± s.e.m. *p < 0.05, two-sided Mann-Whitney test (*n* = 18 WT and *n* = 14 *SRGAP2C* mice). **c**, Relationship between maximum pairwise Euclidean distance of individual smooth vs. R200 trials in neural population state space (as in Fig. 4) and change in performance at each texture difficulty relative to R200 baseline. Each point represents one animal. The grey line shows the pooled regression across all animals. *p < 0.05, Pearson correlation, *n* = 5 WT and *n* = 7 *SRGAP2C* mice.

The parallel observation that *SRGAP2C* mice show both greater neural trajectory separation (**Fig. 4**) and better maintained performance under increased task difficulty (**Fig. 5b**) raises the question of whether these effects are linked at the level of individual animals. To address this, we asked whether the maximum pairwise Euclidean distance between individual smooth and R200 trials in neural state space, measured during 2p imaging when both genotypes performed equivalently on the easy discrimination, predicted the change in performance as texture similarity increased. Across animals, greater peak pairwise separation was associated with less degradation in performance at R1400 (r = 0.61, p = 0.034; **Fig. 5c**), the intermediate difficulty level at which *SRGAP2C* mice show a modest decrease in performance while WT mice show a substantial decline. A trend in the same direction was observed at R800 (r = 0.56, p = 0.061), consistent with representational geometry beginning to predict behavioral capacity at the easier end of the difficulty range. No relationship was observed at R2000 (r = 0.14, p = 0.657), consistent with this texture pair approaching the perceptual limits of both genotypes, where even animals with the greatest neural separation show performance degradation. These results support the view that peak pairwise separation reflects a latent representational capacity that becomes behaviorally limiting when discrimination demands are sufficiently high. Critically, because the neural and behavioral measurements were obtained from independent experimental phases, this relationship cannot be attributed to a within-session performance correlation.

Together, these findings demonstrate that the *SRGAP2C*-mediated change in cortical circuit architecture enhances the transformation of sensory inputs into separable population representations, and that the degree of this separation predicts the capacity to discriminate progressively more similar textures.

### *SRGAP2C* expression restructures directed cortical information flow during task performance

Having established that *SRGAP2C* mice exhibit broader cortical correlation, restructured encoding during motor preparation, enhanced representational geometry, and improved discrimination performance, we next sought to identify the directional structure of the cortical interactions associated with these effects. Specifically, we asked how information flows across the cortical network during task performance.

The enhanced representational geometry and improved discrimination performance observed in *SRGAP2C* mice together point toward a reorganization of how cortical areas interact during task performance. However, correlation and encoding measures cannot resolve whether observed network differences reflect changes in the directional functional influence between regions. This distinction is critical given that *SRGAP2C* enhances long-range cortico-cortical inputs to cortical pyramidal neurons, and that top-down cortical regions show disproportionate encoding increases during motor preparation in *SRGAP2C* mice (**Fig. 3**). Together, this raises the possibility that inter-areal communication is specifically reorganized. To assess whether *SRGAP2C* reshapes the directed influence structure of cortical networks during texture discrimination, we performed Granger causality (GC) analysis^2,44^ on widefield signals across the dorsal cortex (**Fig. 6a**).

**Fig. 6.**
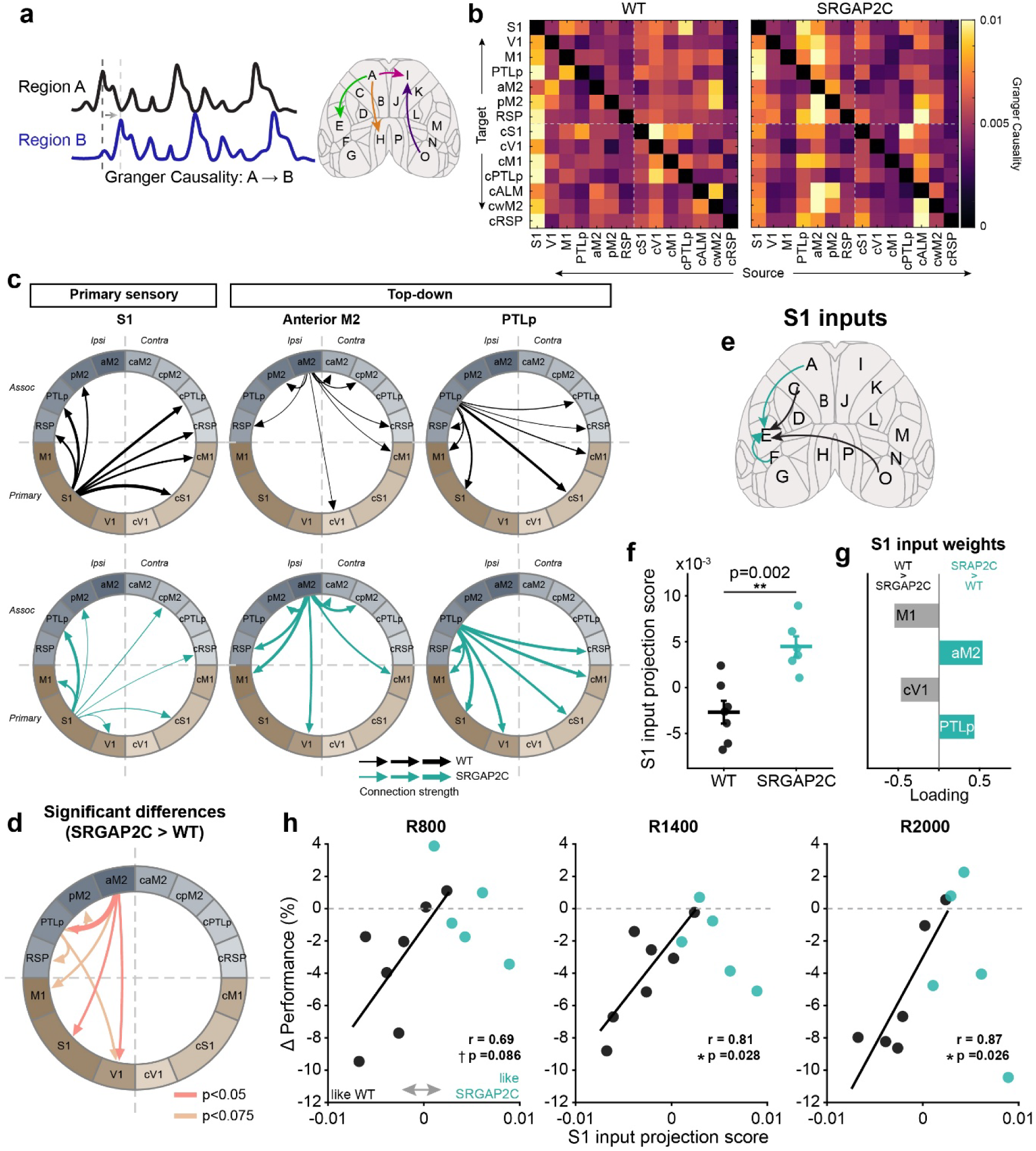
Granger causality analysis reveals a shift toward frontal and associative influence in *SRGAP2C* mice. **a**, Schematic illustrating Granger causality analysis across cortical regions of interest. **b**, Matrices of Granger causality F-statistics for all pairwise region combinations in WT and *SRGAP2C* mice. **c**, Circular node graphs depicting the top 50% connections by Granger causality strength (arrow thickness) for S1, anterior M2, and PTLp in WT (top) and *SRGAP2C* (bottom) mice. **d**, Node graph of statistical differences in connection strength between genotypes. Coral arrows indicate p < 0.05, tan arrows indicate p < 0.075, two-sided Mann-Whitney test, *n* = 7 WT and *n* = 6 *SRGAP2C* mice. **e**, Schematic of dorsal cortex indicating the source regions included in the S1 input pattern analysis. **f**, S1 input projection scores for individual WT and *SRGAP2C* animals. Horizontal lines indicate group means ± s.e.m. Two-sided Mann-Whitney test, **p < 0.01. **g**, Loadings of individual source regions on the S1 input discriminant axis. Positive loadings indicate connections stronger in *SRGAP2C*, negative loadings indicate connections stronger in WT. **h**, Relationship between S1 input projection score and change in psychometric performance relative to R200 baseline at R800, R1400, and R2000 texture difficulties. Each point represents one animal. Solid black lines show linear regression fits for WT animals. Pearson correlation, *p<0.05, *n* = 7 WT and *n* = 6 *SRGAP2C* mice.

GC analysis revealed structured directed connectivity across the dorsal cortex during the preparatory epoch in both genotypes (**Fig. 6b** and **Supplementary Fig. 4**). In WT mice, the strongest directed influences originated from S1, consistent with a feedforward-dominated architecture in which primary somatosensory cortex is the principal driver during task performance (**Fig. 6b, c**). Although the set of regions preferentially targeted by each source area was largely preserved across genotypes, the relative weighting of directed connections was markedly shifted in *SRGAP2C* mice, with aM2 and PTLp contributing a greater share of outgoing influence and S1 contributing comparatively less. This resulted in a net shift in the balance of directed flow away from bottom-up sensory-driven communication and toward top-down associative and frontal sources (**Fig. 6c, d**). Notably, these are the same frontal and associative regions that showed disproportionate encoding increases in the lick-aligned GLM analysis, suggesting that their enhanced directed influence is reflected in their broader task-related recruitment across the cortex.

Taken together, the GC analysis indicates that *SRGAP2C* expression does not rewire the routing architecture of cortical networks but instead shifts the balance of directed influence toward existing top-down pathways, particularly from frontal and associative regions. This rebalancing of directed cortical flow converges with the broader pattern of encoding redistribution and enhanced representational geometry observed across widefield and two-photon analyses, each pointing toward the same reorganization.

The network-level shift toward top-down directed connectivity observed in *SRGAP2C* mice raises a specific question: does this reorganization alter the functional input profile received by S1 neurons in a way that tracks behavioral capacity? This question is particularly motivated by our two-photon imaging results, which established that S1 population trajectory separation is both enhanced in *SRGAP2C* mice and predictive of individual discrimination capacity. Because neural trajectory separation in S1 reflects the integrated influence of all cortical inputs arriving simultaneously rather than the contribution of any single afferent, we reasoned that the behaviorally relevant signal might lie not in any single connection but in the overall composition of inputs to S1. To capture this, rather than testing individual connections in isolation, we characterized the full pattern of directed inputs to S1 as a functional unit.

To do this, we represented the cortical input to S1 from each animal as a vector of Granger causality values across all source regions, capturing the full composition of directed input rather than any single connection. We then identified the axis in this multi-dimensional input space that best separates the WT and *SRGAP2C* group means, i.e., the direction along which the two genotypes differ most consistently in their overall S1 input composition. Projecting each animal onto this axis yields a scalar score reflecting the degree to which their S1 input profile resembles the *SRGAP2C* configuration relative to WT. However, because this projection is computed from the full dataset, individual source regions may contribute to the separation by chance. To identify which regions reliably distinguish the two genotypes, we used a leave-one-out procedure: we recomputed the discriminant axis iteratively, each time excluding one animal, and retained only regions whose contribution was consistent in direction across all subsamples. This conservative stability criterion ensures that the identified sources reflect a genuine and replicable feature of the genotype difference rather than the influence of any single animal. This confirmed that the S1 input pattern is dominated by two opposing contributions: M1 and contralateral V1 contribute disproportionately to S1 in WT mice, while aM2 and PTLp contribute disproportionately in *SRGAP2C* mice (**Fig. 6e-g**). This pattern is consistent with the network-level shift described above and provides connection-level specificity for the top-down reorganization.

The S1 input projection scores differed significantly between genotypes (**Fig. 6f**), confirming that the overall composition of functional cortical input to S1 is systematically shifted in *SRGAP2C* mice. Because this reorganization is expressed as a shift in the relative balance of multiple cortical inputs to S1 rather than a change confined to any single projection, we next asked how it relates to behavioral performance. Critically, within WT animals, those whose S1 input profile more closely resembled the *SRGAP2C* configuration (higher projection score values) showed less degradation in discrimination performance as texture similarity increased, with significant correlations at intermediate and high difficulty (R1400: r = 0.81, p = 0.028; R2000: r = 0.87, p = 0.026; **Fig. 6h**) and a consistent trend at the easiest step above baseline (R800: r = 0.69, p = 0.086). Within *SRGAP2C* mice, no significant relationship was observed between input projection score and discrimination performance, potentially reflecting a ceiling in input reorganization within this group.

Together, these results show that *SRGAP2C* does not simply strengthen individual top-down connections to S1 but shifts the overall balance of its cortical input profile toward association and frontal sources. Within a single genotype, this shift in input composition tracks individual discrimination capacity, converging with the independent population-geometry result to suggest that directed input reorganization and representational separability reflect the same underlying network reconfiguration.

## DISCUSSION

In this study, we investigated how expression of the human-specific gene *SRGAP2C* shapes cortical network dynamics and sensorimotor learning during a sensory discrimination task. By combining widefield and two-photon calcium imaging, generalized linear modeling, directed connectivity analysis, and psychometric behavioral testing, we traced an arc from network-level organization to population-level representational geometry to behavioral performance. Our central findings show that *SRGAP2C* reshapes cortical network organization by shifting the balance of directed influence toward top-down frontal and associative sources, that this rebalancing is reflected in the composition of cortical input onto S1, and that it is accompanied by more separable population representations in S1 during motor preparation. Moreover, across independent measures, the degree of this reorganization tracked discrimination performance specifically as task demands increased, linking the network-level changes to behavior.

Across both the stimulus and preparatory epochs, a consistent pattern emerged where *SRGAP2C* mice showed broader spatial distribution of correlated and task-relevant activity, while the total amount of task-relevant information captured by the network remained equivalent to WT. During texture contact, the expanded co-activation seeded from somatosensory cortex likely reflects increased propagation of sensory representations across the cortical surface, amplified by enhanced cortico-cortical wiring, without reorganization of their structure. Population representations in S1 during this epoch are primarily shaped by feedforward thalamocortical input^23,45^, which is equivalent between genotypes, and the broader correlation pattern in *SRGAP2C* mice reflects increased spread of these representations rather than a change in their content.

The functional consequences of encoding redistribution become most apparent during the preparatory epoch preceding lick execution, during which we observed distributed encoding across the cortex of *SRGAP2C* mice, greater Euclidean distance between texture trajectories in somatosensory population state space, and enhanced directed influence from frontal cortex to a broad network of cortical targets. These independent lines of evidence converge on the same reorganization, centered on a coordinated reweighting of cortical input to S1: PTLp and aM2 emerge across the encoding, directed connectivity, and input-composition analyses as disproportionately recruited in *SRGAP2C* mice, while the input-composition analysis additionally identifies M1 and contralateral V1 as sources that contribute disproportionately in WT. This four-source reweighting, frontal and associative sources up, primary motor and visual sources down, is consistent with a model in which top-down signals from higher-order areas, the same projections enhanced by *SRGAP2C*, project back onto sensory cortex and shape population activity during the period when the brain transforms sensory information into a motor plan. The enhanced cortico-cortical connectivity in *SRGAP2C* mice thus appears to shape representational geometry specifically during this feedback-dominated epoch, increasing the separation between population trajectories. Critically, this top-down broadcast supplements rather than replaces the conserved sensorimotor transformation pathway: the fundamental routing architecture is preserved across genotypes, but its relative weighting is shifted, layering additional preparatory signaling onto the feedforward scaffold. The net result is a redistribution of where task-relevant encoding is expressed across the cortex, without a change in how much task-relevant variance the network captures overall, i.e., the same information is encoded, but through a more spatially distributed network organization.

Beyond the network-level rebalancing, the composition of cortical input onto S1 provides connection-level specificity for this reorganization: the shift is not an abstract redistribution of directed flow but a measurable change in what S1 actually receives, with PTLp and aM2 contributing disproportionately in *SRGAP2C* mice and M1 and contralateral V1 dominating in WT. The functional relevance of this change is supported by variation within WT animals, such that animals whose input profile more closely resembled the *SRGAP2C* configuration maintained performance better as textures became more similar. Because this relationship holds within a single genotype, it cannot solely be attributed to the between-group differences that accompany *SRGAP2C* expression. Notably, the input-profile metric predicted performance at the most demanding texture (R2000) where peak trajectory separation did not, indicating that the two readouts capture related but non-identical aspects of the reorganization: trajectory separation reflects how far population representations are driven apart, whereas input composition may carry additional behaviorally-relevant structure not reduced to the distance between trajectory means.

The behavioral consequences of this reorganization became apparent when discrimination demands increased. On the standard rough versus smooth task, both genotypes performed equivalently, consistent with the interpretation that feedforward sensory processing is sufficient for simple, high-contrast discrimination. When texture similarity was parametrically increased however, *SRGAP2C* mice maintained performance at difficulty levels where WT mice showed substantial deficits, consistent with the state space analysis. The individual-animal correlation between peak pairwise trajectory separation and psychometric performance directly connects these neural and behavioral observations. At R200, the discrimination is tractable for both genotypes and behavioral variance is limited, while at R800 and R1400, individual differences in representational geometry become the binding constraint. At R2000, the task exceeds the capacity of both genotypes and the relationship breaks down. The broader implication is that *SRGAP2C* does not simply shift mean performance upward, but rather expands the representational capacity of individual animals in a way that is behaviorally expressed only when discrimination demands approach the limits of what the circuit can reliably support.

One notable aspect of our findings is that the expanded S1-seeded connectivity in *SRGAP2C* mice emerged with task expertise rather than being present from the outset of training, ruling out the possibility that *SRGAP2C*-mediated connectivity passively increases activity spread regardless of behavioral context. Instead, this result suggests that learning actively recruits the expanded cortico-cortical substrate afforded by *SRGAP2C* expression. This would be consistent with the broader principle that cortical circuits are shaped by the interaction between structural connectivity and behavioral experience, and raises the possibility that *SRGAP2C* expands the repertoire of network configurations available to the cortex during learning rather than imposing a fixed pattern of altered dynamics, a question that warrants systematic investigation across training stages in future work.

Because *SRGAP2C* is expressed from early development, we cannot fully dissociate altered ongoing circuit function from connectivity established developmentally. Beyond this, *SRGAP2C* does not act within a single pathway but coordinately reshapes the relative functional balance of inputs to S1, with aM2 and PTLp contributing proportionally more and M1 and contralateral V1 proportionally less, in a task-dependent manner. Because this effect is expressed as opposing changes across multiple sources, it cannot be isolated by perturbing any single projection: removing one arm of the reweighting reconstructs the configuration of neither genotype but instead produces a state present in neither. Suppressing aM2 or PTLp, for instance, would shift the input balance opposite to the *SRGAP2C* signature while leaving the coordinated reweighting itself untested. Resolving how this distributed reorganization shapes S1 geometry, directly, through local inhibition, or via multisynaptic routes^46^, will require combinatorial, multi-site approaches. In addition, beyond the dorsal cortical surface accessible to widefield imaging, *SRGAP2C*-mediated connectivity changes likely extend to deeper structures and subcortical circuits. *SRGAP2C* was previously shown not to increase thalamocortical inputs, indicating that the effects observed here are driven primarily by cortico-cortical interactions. However, whether enhanced cortical output alters the engagement of downstream subcortical targets remains open.

*SRGAP2C* arose as a human-specific gene duplication during a period of neocortical expansion in the *Homo* lineage marked by a pronounced increase in the density and reach of cortico-cortical projections. Previous work established that *SRGAP2C* increases cortico-cortical connectivity, but the functional consequences for cortical computation underlying behavior had remained unclear. Our findings indicate that *SRGAP2C* expression restructures how cortical networks facilitate the transformation of sensory evidence into motor preparation by shifting the balance of cortical coordination toward top-down frontal sources across a broader cortical network, yielding more separable population representations and improved discrimination under demanding conditions. Because these conclusions derive from a humanized mouse model, they isolate the component of cortical reorganization attributable to *SRGAP2C* rather than reconstructing the integrated effects of the many genetic and architectural changes that distinguish the human cortex. Establishing the generality of these network-level principles will require cross-species comparison and validation in human cellular models. With that scope in mind, these results suggest that the evolutionary elaboration of cortico-cortical connectivity may have expanded cognitive capacity not by adding new sensory machinery, but by enhancing the brain’s ability to coordinate how sensory evidence is transformed into action.

## METHODS

### Mice

All animal procedures were done according to protocols approved by the Institutional Animal Care and Use Committee (IACUC) at the Medical University of South Carolina. Adult male and female mice (>P65) were used for all experiments. Conditional *SRGAP2C*-expressing mice heterozygous for the *Rosa26*^LSL-SRGAP2C^ knock-in allele, as previously described^18^, were originally generated in the Polleux laboratory at Columbia University and maintained as an in-house colony. In this study, we refer to these mice as *SRGAP2C* mice. For widefield calcium imaging experiments, these mice were crossed with Thy1-GCaMP6f transgenic mice (Jackson Labs; C57BL/6J-Tg(Thy1-GCaMP6f)GP5.17Dkim/J). For two-photon calcium imaging experiments, mice received intracortical injections to achieve pan-neuronal expression of the calcium indicator jGCaMP8m^47^. Mice were allowed 2 weeks post-injection for sufficient expression prior to imaging. All lines were maintained on a C57BL/6J background. Mice were weaned at postnatal day 28 and housed with littermates in temperature and humidity-controlled facilities with a 12-hour light/dark cycle and *ad libitum* access to food and water.

### Behavioral task

#### Behavioral setup

Behavioral experiments were conducted in a custom-built black enclosure with a light-blocking door. The behavioral apparatus was adapted from a previously described system^18,19^. A stepper motor (bipolar, 200 steps/rev, 28×32mm, 3.8V, 0.67 A/Phase; Pololu #1205) rotated laser-cut acrylic textures into position, and a linear actuator (Actuonix L12-30-50-6-R) advanced the selected texture to within ∼1 cm of the right whisker pad. Laser-cut acrylic textures measured 16 × 33 mm and contained laser-cut vertical grooves approximately 500 µm deep, 350 µm wide, and spaced 200 µm apart (or 800 µm, 1400 µm, and 2000 µm for the psychometric curve experiment). The cut side was presented as the rough texture and the uncut side was used as the smooth texture. A fan (Cooler Master 80 mm Silent Fan) directed airflow away from the mouse to minimize the use of olfactory cues, and a white-noise generator masked ambient sounds. Water rewards (∼5 µl) were delivered through stainless-steel lick ports via a solenoid valve (The Lee Company, LFAA1209512H) and licking was detected using capacitive touch sensors (Sparkfun MPR121). Task control was implemented using an Arduino Uno and task-relevant signals were logged in MATLAB. For imaging sessions, behavioral signals were simultaneously recorded through the same data-acquisition board (National Instruments) used for the imaging systems and aligned post hoc to the neural data.

#### Behavior training and testing

Behavioral training and testing procedures were adapted from a previously described protocol^18,19^. Mice were denied water access in the home cage and learned to receive water during behavioral training. During the first week, mice were habituated to head fixation and familiarized with the lick ports and water reward delivery system. Following habituation, mice underwent one week of forced alternation training, in which rough and smooth textures were presented in strict alternation, allowing mice to learn the trial structure and texture-port associations. Mice were trained to report texture identity by licking the left port for rough texture presentation and the right port for smooth texture presentation. Following texture presentation and retraction, a brief delay period was enforced, after which a light cue signaled the opening of the response window. Auditory tones indicated trial outcomes and the start of the subsequent trial (low-frequency tone for trial completion; high-frequency tone for aborted trials due to early licking). Incorrect trials were punished with a 10 sec timeout prior to the subsequent trial. If no lick responses were detected within 30 seconds, the trial timed out followed by the start of the next trial. Following forced alternation training, mice were tested on the full two-alternative forced choice (2AFC) task, in which rough and smooth textures were presented in random order. All other task parameters remained identical to those used during forced alternation training. Performance was calculated as the fraction of correct trials completed within a session.

#### Psychometric curve analysis

Following expert training on the rough (R200) versus smooth texture discrimination, mice underwent psychometric curve testing to assess discrimination performance across a range of texture similarities. The standard rough texture (R200, 200 µm groove spacing) was replaced with progressively smoother textures (R800, R1400, and R2000), while the smooth texture remained unchanged. Each texture was cleaned with isopropyl alcohol before use and tested over three consecutive sessions to accumulate sufficient trials for analysis while minimizing the potential for mice to learn the harder discriminations across sessions. Trial counts and task parameters were otherwise identical to those used during expert testing. To confirm that mice were using texture as the discriminative cue rather than incidental non-textural cues, a smooth versus smooth control session was conducted following completion of the full psychometric curve. Mice that maintained expert performance were considered to be relying on non-textural cues (‘cheaters’) and were excluded from all analyses. For each mouse, performance at each texture was calculated as the fraction of correct trials per session, averaged across the three sessions. To account for individual differences in baseline expert performance, the performance of each mouse at each texture was normalized by expressing it as a change relative to their mean expert performance on the standard rough versus smooth discrimination, yielding a measure of performance drop as texture difficulty increased. Group-level psychometric curves were constructed by averaging normalized performance values across mice within each genotype.

### Widefield imaging

#### Surgical procedure

Mice were anesthetized with isoflurane (5% induction and 2% maintenance) and secured in a stereotaxic frame (Neurostar). Meloxicam (5 mg/kg body weight) was injected subcutaneously and lidocaine was injected at the incision site. An incision along the midline of the head was made and the skin was retracted to expose the dorsal skull. Connective tissue was carefully cleared away and the scalp margins were affixed to the skull using VetBond. The skull surface was cleared by applying three thin layers of cyanoacrylate glue (Zap-A-Gap CA+, Pacer technology), allowing each layer to polymerize before the next layer was added. A custom-cut titanium headplate (MPFI Mechanical Workshop) was positioned over the dorsal cortex and secured to the skull using dental acrylic (C&B Metabond, Parkell) to enable stable head fixation during imaging and behavioral training. After completion of all surgical preparations, mice recovered on a heated pad and were returned to their home cage once fully ambulatory.

#### Image acquisition

Widefield calcium imaging of GCaMP fluorescence was performed during behavioral task engagement using a dual-wavelength strobed illumination system^27,28^. The exposed dorsal cortex was illuminated sequentially with a blue LED (Thorlabs M470L5) passed through a 460/60 nm band-pass filter (Semrock FF01-460/60-25) to excite GCaMP, and a green LED (Thorlabs M530L3) passed through a 530/43 nm band-pass filter (Semrock FF01-530/43-25) to measure total hemoglobin absorption. A 565/133 nm band-pass emission filter (Semrock FF01-565/133-25) was placed in front of the camera to block excitation light. Near-simultaneous acquisition of GCaMP and hemodynamic signals enabled correction of wavelength-dependent absorption changes known to confound widefield calcium data^27,28^. Fluorescence was recorded using an Andor Zyla 4.2+ sCMOS camera coupled to a Nikon Micro-NIKKOR 60 mm f/2.8D lens at 60 Hz (30 Hz per LED channel). Imaging was performed continuously for 1 hour while mice performed the behavioral task. Signals from the imaging and behavior systems were acquired through a shared data-acquisition board (National Instruments) and aligned post-hoc during preprocessing.

#### Image preprocessing and analysis

All image processing and analyses were performed in MATLAB (MathWorks). Images were separated into GCaMP fluorescence (blue channel) and hemodynamic absorption (green channel) frames based on LED strobe timing. To reduce high-frequency noise in the hemodynamic signal, the green channel was low-pass filtered at 2 Hz prior to hemodynamic correction. Hemodynamic correction was then applied using the approximate division method for single-wavelength reflectance data, as described previously^27^, in which the GCaMP fluorescence signal is normalized by the low-pass filtered green channel reflectance to remove wavelength-dependent absorption changes caused by fluctuations in total hemoglobin. Slow signal drifts were subsequently removed by subtracting a slowly-varying baseline estimated from this ΔF/F signal.

#### Correlation and Latency Analysis

Prior to analysis, all widefield images were registered to the Allen Common Coordinate Framework (CCF) using the registration pipeline from LocaNMF^48^. ROIs were defined as square regions expanded from the centroid of each Allen CCF-defined area. Secondary motor cortex (M2) was subdivided into an anterior and a posterior ROI based on the CCF boundary.

Functional connectivity was assessed by computing Pearson correlation coefficients between the mean ΔF/F time series of a seed ROI in primary somatosensory cortex (S1) and all other cortical pixels, generating seed-based correlation maps for each session. Correlation was computed within the peri-stimulus time window (±1 sec around full texture contact). Epochs surrounding lick events were excluded from correlation analysis, as movement artifacts during licking produce widespread correlated activity across the cortex that precludes meaningful interpretation of functional connectivity.

#### Generalized linear model analysis

To quantify the relationship between widefield GCaMP fluorescence and task and behavioral variables, we constructed a linear ridge regression encoding model, as previously described^30,31^. Dimensionality reduction was first applied to the GCaMP data using singular value decomposition (SVD), retaining the top components that collectively explained >95% of the variance (∼200 components). A design matrix was constructed using task-related regressors aligned to either texture contact or lick onset, along with continuous whisker motion traces and binary body movement bout regressors derived from infrared video recordings (Teledyne FLIR Blackfly S USB3 [model # BFS-U3-04S2M-CS: 0.4 MP, 522 FPS, Sony IMX287, Mono]). Movement bouts were expanded by 1 second before onset and 2 seconds after offset, as described previously^31^. All regressors were z-scored prior to model fitting.

The model was fit using task and movement regressors combined. For each SVD component, the regularization parameter (λ) was optimized by minimizing the negative marginal log-likelihood of the residuals using a quasi-Newton algorithm. Model performance was evaluated using 5-fold cross-validation to obtain unbiased estimates of variance explained (R²). Beta weights from a final model refit on all data were projected back into pixel space by multiplying with the SVD spatial components, yielding whole-brain maps of regressor-specific activity.

#### Granger Causality Analysis

Directed functional connectivity between cortical ROIs was assessed using pairwise conditional Granger causality (GC), implemented using the MVGC Multivariate Granger Causality Toolbox^44^. All analyses were performed on lick-aligned trial epochs spanning approximately 3.5 to 0.5 seconds before lick onset, chosen to capture task-relevant neural dynamics while avoiding movement artifacts associated with licking, as described in the main text. ROI time series were extracted as the mean ΔF/F signal within each region and preprocessed for stationarity prior to GC estimation. Each trial time series was linearly detrended and mean-subtracted, then low-pass filtered at 5 Hz. The optimal autoregressive model order was determined independently for each animal using the Akaike Information Criterion (AIC), with a maximum model order of 20 lags. VAR models were estimated using the Levinson-Wiggins-Robinson (LWR) algorithm, and model stationarity was confirmed by verifying that the spectral radius of the companion matrix was less than 1. GC F-statistics were computed for each directed ROI pair.

#### S1 cortical input profile analysis

To characterize the overall pattern of directed cortical inputs to S1 as a functional unit, we extracted the GC F-statistics from all source regions to S1 for each animal, forming a vector representation of the S1 input profile of each animal. A mean-difference discriminant projection was computed by subtracting the WT group mean input vector from the *SRGAP2C* group mean input vector and normalizing the resulting difference vector to unit length. Each animal received a scalar projection score equal to the dot product of their S1 input vector with this unit projection axis, reflecting the degree to which their S1 input profile resembled the *SRGAP2C* configuration relative to WT. To identify which source regions stably contributed to the genotype separation, we performed leave-one-out (LOO) analysis by iteratively excluding each animal and recomputing the discriminant projection on the remaining animals. For each source region, we computed the mean absolute loading magnitude across all LOO iterations and the sign consistency. Regions were retained for the final discriminant if their mean absolute loading exceeded the 70th percentile across all source regions and their sign consistency was at least 80%. The projection axis used for all reported analyses was computed on the full dataset using only these stably contributing regions.

### Two-photon imaging

#### Surgical procedure

Mice were anesthetized with isoflurane (5% induction and 2% maintenance) and secured in a stereotaxic frame (Neurostar). Meloxicam (5 mg/kg body weight) was injected subcutaneously and lidocaine was injected at the incision site. A full-thickness circular craniotomy was made over the barrel field of primary somatosensory cortex (centered at 1.5mm caudal to bregma and 3mm lateral to the midline) using a high-speed drill, and the bone flap was removed. A Robot Stereotaxic system (Neurostar) was used to identify the injection site and deliver AAV1-syn-jGCaMP8m-WPRE (Addgene viral prep #162381-AAV1; ^47^). Following injection, a pre-assembled double-layer glass window, consisting of a 3mm coverslip bonded to a 5mm coverslip (using Norland Optical Adhesive NOA 78, Edmund Optics), was placed over the craniotomy with only the 3mm coverslip seated against the dura. A custom-cut titanium headplate (MPFI Mechanical Workshop) was then positioned around the window and secured to the skull using dental acrylic (C&B Metabond, Parkell) to allow for stable head fixation during imaging and behavioral training. After completion of all surgical preparations, mice recovered on a heated pad and were returned to their home cage once fully ambulatory. Mice were allowed 2 weeks post-injection and craniotomy for sufficient expression prior to imaging.

#### Image acquisition and data preprocessing

Two-photon calcium imaging was performed using a Bergamo II microscope (Thorlabs) running ThorImage LS software, with hardware synchronization via ThorSync. Excitation light was provided by a tunable femtosecond pulsed laser (Coherent Chameleon Discovery NX with Total Power Control) set to 940 nm. Fluorescence was collected through a 16× 0.8 NA objective (Nikon) at approximately 30 Hz. Images were acquired at 512 × 512 pixels covering a 422 × 422 µm field of view, at cortical depths of 150-250 µm below the pial surface, corresponding to layer 2/3.

Non-rigid motion correction of acquired images was performed using NoRMCorre^49^. Neuronal ROIs were manually identified using the Cell Magic Wand tool (Fitzpatrick Lab, Max Planck Florida Institute for Neuroscience) in ImageJ and time-courses were extracted as the mean fluorescence across all pixels within each ROI. Neuropil contamination was estimated and removed as previously described^18^. Only animals in which >100 neurons were successfully identified in the field of view were included in subsequent analyses to ensure that trajectory analyses were performed on populations of sufficient size to yield stable low-dimensional representations. Timeout trials and trials flagged as forced were excluded from all analyses. For each neuron and trial, ΔF/F was computed where baseline was defined as the mean fluorescence during the 0.5 seconds immediately preceding texture extension. Individual trial traces were subsequently smoothed using MATLAB’s smoothdata function.

#### Neural trajectory analysis

Population-level neural trajectories were analyzed in a reduced-dimensional state space to characterize the geometry of stimulus representations. Analyses were performed on neuropil-corrected ΔF/F traces and restricted to neurons exhibiting a mean positive response during the task epoch, defined as a mean ΔF/F greater than zero across all trials and timepoints, focusing the geometric analysis on the subpopulation that carries positive stimulus-driven signal. Trial counts were balanced across stimulus conditions (rough vs. smooth) by subsampling to the minimum number of trials available across both conditions for each animal.

Principal component analysis (PCA) was applied to the full single-trial dataset with both conditions concatenated and the top 10 PCs were retained. Individual trials were then projected into this common PC space. Prior to distance computations, each neuron’s mean activity during baseline frames (1-15) was subtracted from all timepoints to re-zero trajectories at trial onset, isolating stimulus-driven dynamics from baseline activity.

Stimulus separability was quantified using two complementary distance metrics computed in PC space. The Euclidean distance between condition means was calculated at each timepoint as the distance between the mean trajectory of rough trials and that of smooth trials. Pairwise between-condition distances were additionally computed across all cross-condition trial pairs to characterize trial-level separability. To confirm that observed distances were not attributable to chance, a shuffled null distribution was generated for each animal by pooling trials across conditions, randomly splitting them into two equal halves in PCA space across 200 iterations, and computing the distance between split-half mean trajectories at each timepoint.

To examine whether individual differences in neural trajectory separation predict psychometric discrimination capacity, the maximum pairwise Euclidean distance between individual smooth and R200 trials was computed per animal as a summary measure of peak representational separability. This maximum was taken across all cross-condition trial pairs and all timepoints within the trial epoch. Psychometric performance was quantified as delta performance: the difference between each animal’s median proportion correct across sessions at each texture difficulty (R800, R1400, R2000) and their median performance on the standard R200 discrimination. The relationship between peak pairwise separation and delta performance was assessed using Pearson correlation across all animals.

#### Statistics and reproducibility

Statistical analysis was performed using MATLAB (MathWorks). Group differences were assessed using the Mann-Whitney U test (Wilcoxon rank-sum). Differences in learning rate between genotypes were assessed using linear regression with a genotype x session interaction term. Distributional differences were evaluated using the two-sample Kolmogorov-Smirnov (KS) test. For Granger causality analyses, statistical significance was assessed using Granger’s F-test with false discovery rate (FDR) correction for multiple comparisons (Benjamini-Hochberg procedure, α = 0.05), while for group-level comparisons between WT and SRGAP2C mice, mean GC F-statistics for each directed connection were compared using two-sided Wilcoxon rank-sum tests, with effect sizes reported as rank-biserial correlation coefficients. A test was considered significant when *P*<0.05. Littermate controls were drawn from the same crosses and were negative for either the NexCre driver or the Rosa26^LSL-SRGAP2C^ allele (while carrying the other), ensuring that genotype comparisons control for both Cre expression and the presence of the knock-in locus. For wide-field imaging, 6 *SRGAP2C* mice and 7 littermate controls, and for two-photon imaging, 7 *SRGAP2C* and 5 littermate controls, were obtained from a minimum of four independent litters. For behavioral analyses, a total of 16 *SRGAP2C* and 19 WT littermate controls were obtained from 12 independent litters. Of these, 14 *SRGAP2C* and 18 WT mice completed psychometric curve testing following expert training. Independent data points shown denote data from individual animals.

## ACKNOWLEDGEMENTS

We thank Dr. Arjun Bharioke and Dr. Takashi Sato for insightful comments, experimental guidance, and feedback on the manuscript. We thank all members of the Schmidt lab for valuable discussions and input. This work was supported by NIH R00 (R00 NS109323) (ERES) and a Whitehall Foundation Research Grant (2022-05-035) (ERES).

## AUTHOR CONTRIBUTIONS

Conceptualization: HTZ, ERES. Methodology: HTZ, ERES. Surgeries: HTZ. Behavioral training: HTZ, TA. Widefield imaging: HTZ. Two-photon imaging: HTZ, TA. Data Analysis: HTZ, ERES. Funding acquisition: ERES. Writing- original draft: HTZ, ERES.

## SUPPLEMENTARY FIGURES

**Supplementary Fig. 1.**
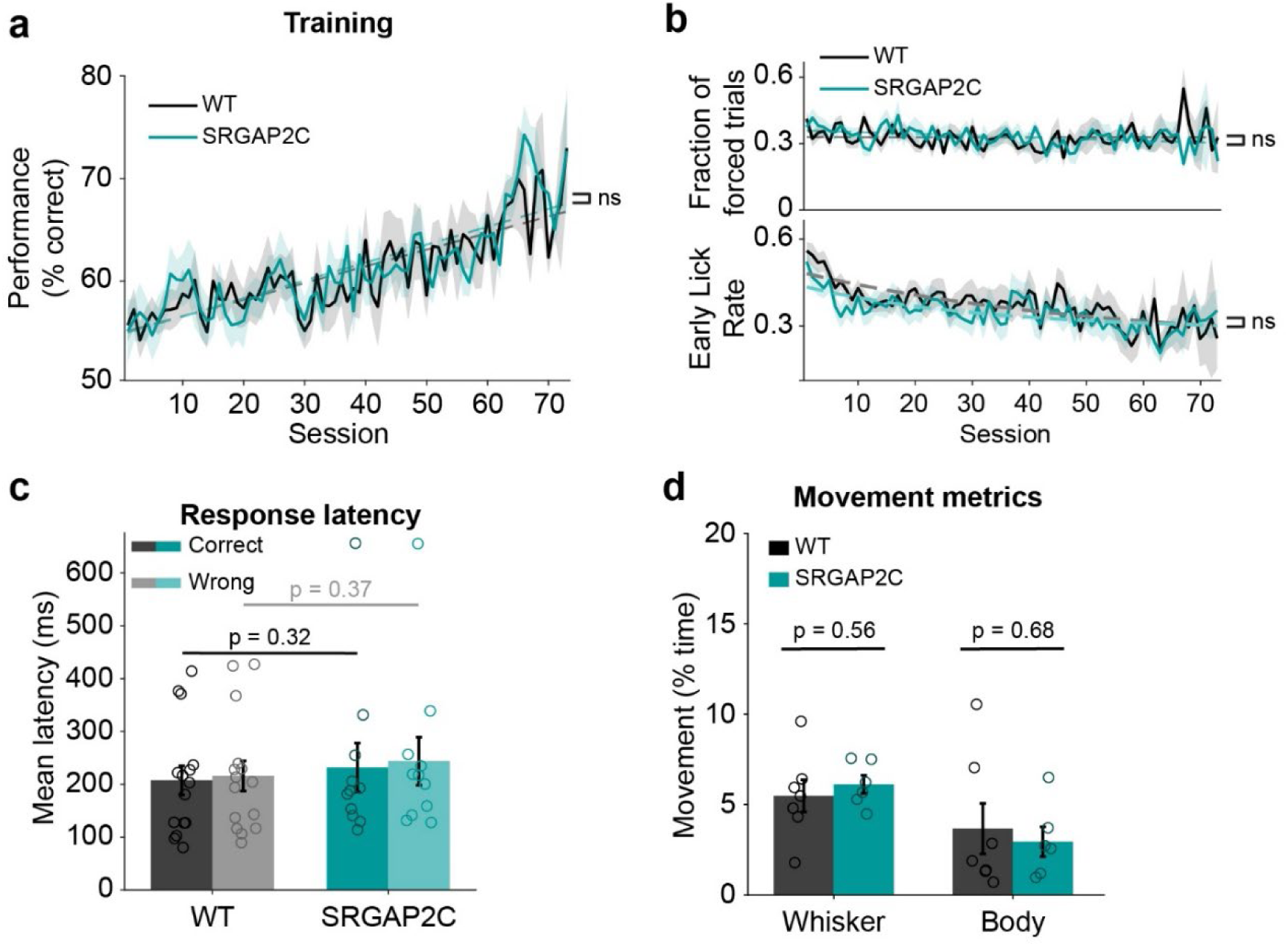
WT and *SRGAP2C* mice show comparable behavioral performance and movement profiles. **a**, Discrimination performance across training sessions for WT and *SRGAP2C* mice, shown as mean with shaded areas indicating ± s.e.m. (ns, linear regression). **b**, Fraction of forced trials per session (top) and early lick rate per session (bottom), shown as mean with shaded area indicating ± s.e.m (ns, linear regression). **c**, Response latency for correct and wrong trials. p values shown for all pairwise comparisons. n = 19 WT, n = 16 *SRGAP2C* mice (a-c). **d**, Whisker movement and body movement, each expressed as percentage of time active during the trial period. Bar graphs plotted as mean ± s.e.m. Open circles indicate individual animals (*n* = 7 WT and *n* = 6 *SRGAP2C* mice).

**Supplementary Fig. 2.**
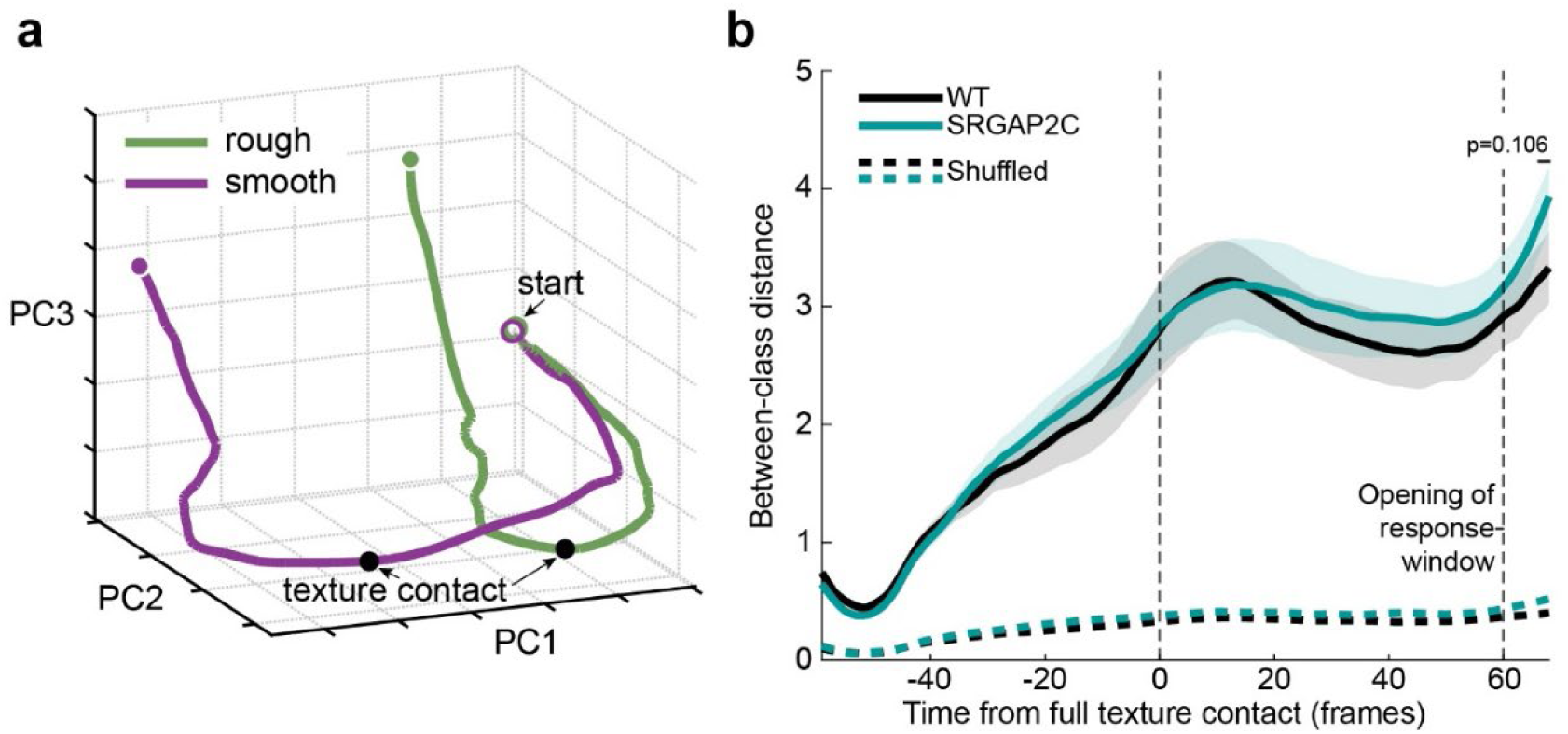
Separation of neural trajectories for texture-aligned trials. **a**, Texture-aligned neural trajectories projected onto the top three principal components (PC1-PC3) for smooth and rough trials in a representative animal. Black dots indicate the moment of full texture contact. **b**, Euclidean distance between rough and smooth neural trajectories for WT and *SRGAP2C* mice as a function of time relative to full texture contact, shown as mean with shaded area indicating ± s.e.m., compared to shuffled null distributions (dashed lines). First vertical dashed line indicates full texture contact, second vertical dashed line indicates opening of the response window, two-sided Mann-Whitney test (*n* = 5 WT and *n* = 7 *SRGAP2C* mice).

**Supplementary Fig. 3.**
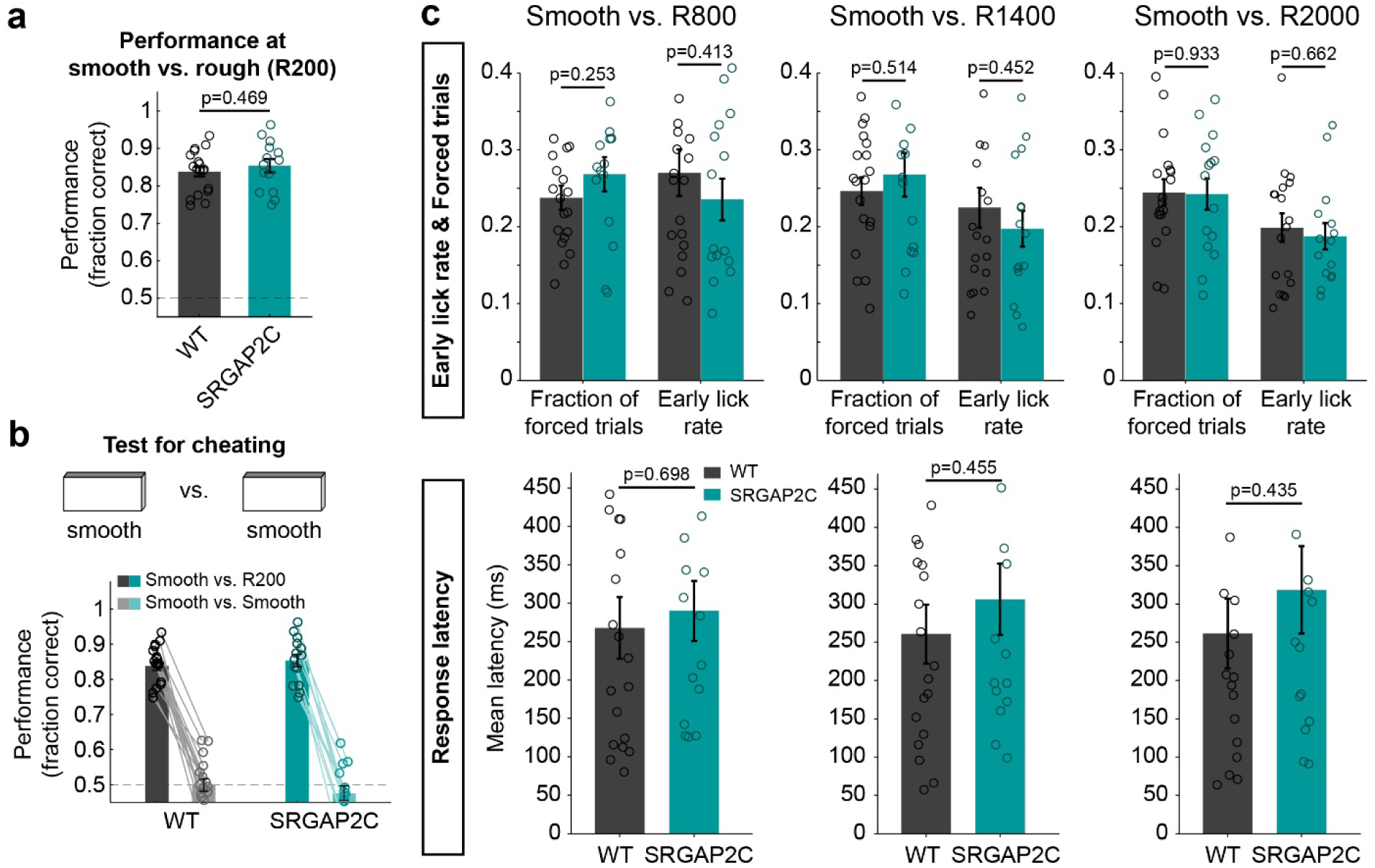
Performance metrics across texture and control conditions for WT and *SRGAP2C* mice. **a**, Discrimination performance at smooth vs R200 for WT and *SRGAP2C* mice. **b**, Bar graph showing discrimination performance on a smooth vs. smooth control session following psychometric curve testing, for WT and *SRGAP2C* mice. Dashed line indicates chance performance (50%). **c**, Performance metrics for WT and *SRGAP2C* mice for each texture (R800, R1400, and R2000). Two-sided Mann-Whitney test, bar graphs plotted as mean ± s.e.m. Open circles indicate individual animals (*n* = 18 WT and *n* = 14 *SRGAP2C* mice).

**Supplementary Fig. 4.**
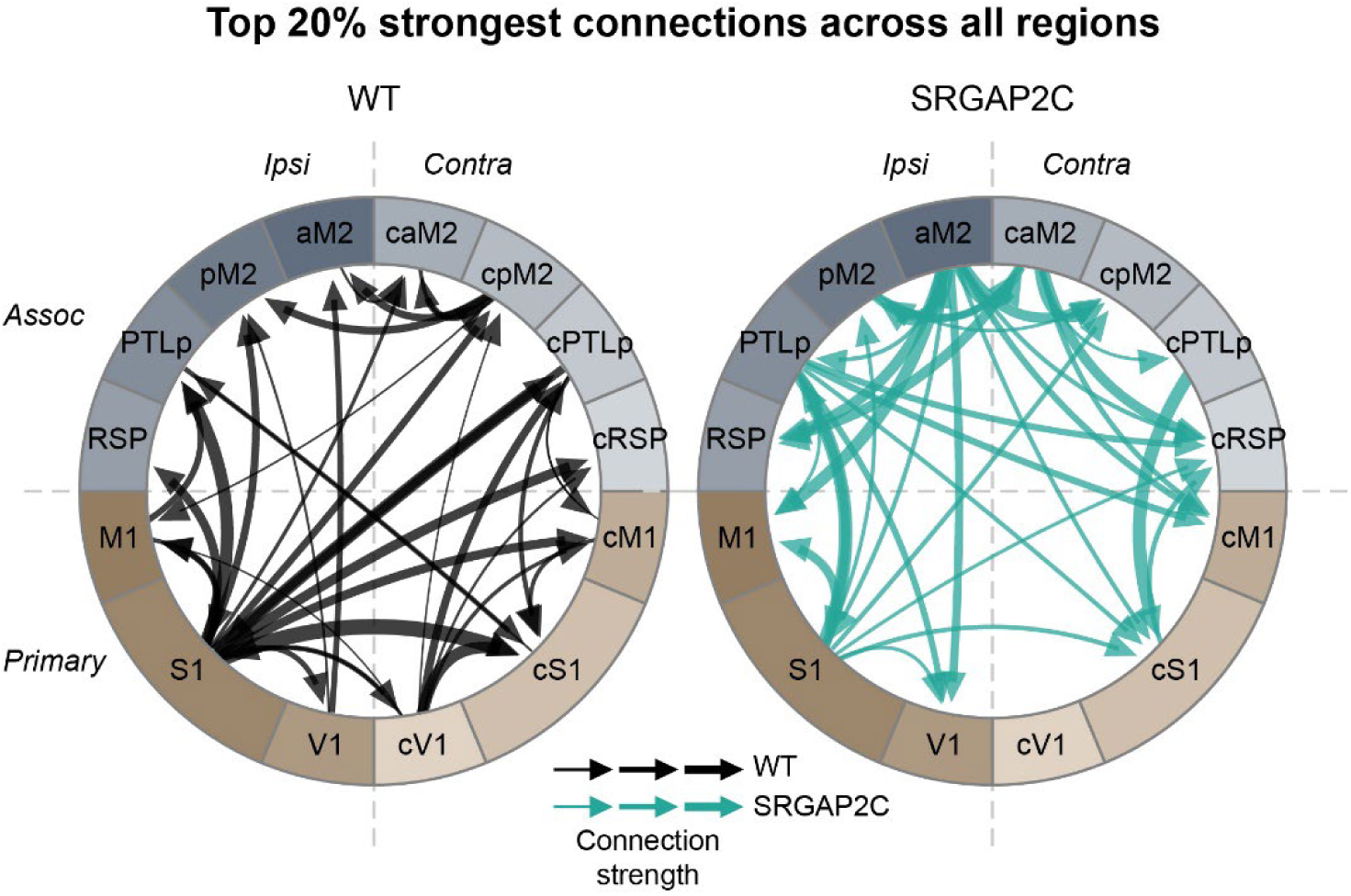
Cortex-wide Granger causality connectivity during the lick-aligned epoch. Node graphs depict the top 20% of directed connections by Granger causality strength across all cortical regions of interest for WT (left) and *SRGAP2C* (right) mice, with arrow thickness proportional to connection strength (*n* = 7 WT and *n* = 6 *SRGAP2C* mice).

